# Requirement of the Dynein-adaptor Spindly for mitotic and post-mitotic functions in *Drosophila*

**DOI:** 10.1101/199414

**Authors:** Giuliana D. Clemente, Matthew R. Hannaford, Jens Januschke, Eric R. Griffis, Hans-Arno J. Muller

## Abstract

Spindly is a mitotic checkpoint protein originally identified as a specific regulator of Dynein activity at the kinetochore. In metaphase, Spindly recruits the Dynein/Dynactin complex, promoting the establishment of stable kinetochore-microtubule interactions and progression into anaphase. While details of Spindly function in mitosis have been worked out in cultured human cells and in the *C. elegans* zygote, the function of Spindly within the context of an organism has not yet been addressed. Here we present loss- and gain-of-function studies of Spindly in *Drosophila*. We investigated the requirements of distinct protein domains for the localisation and function of Spindly. We find that knock-down of Spindly results in a range of mitotic defects in the female germ line and during cleavage divisions in embryogenesis. Overexpression of Spindly in the female germ line is embryonic lethal and results in altered egg morphology. To determine whether Spindly plays a role in post-mitotic cells we altered Spindly protein levels in migrating cells and found that ovarian border cell migration is sensitive to the levels of Spindly protein. Our study uncovers novel functions of the mitotic checkpoint protein Spindly in *Drosophila*.

## Introduction

Faithful chromosome segregation in mitosis is fundamental for maintaining genome stability and cell viability. Segregation of chromosomes relies on the mitotic spindle, a bipolar microtubule-based machinery that assembles at prometaphase of mitosis. The mitotic spindle needs to unequivocally recognise and bind on each sister chromatids a multi-protein complex called kinetochore that provides a unique end-on attachment site for microtubules. The presence of misaligned chromosomes activates the Spindle Assembly Checkpoint (SAC), which prevents anaphase until each chromosome is bound to the spindle. The SAC is activated via Aurora B and Mps1-dependent assembly of a mitotic checkpoint complex (MCC) which inhibits the activity of the Anaphase Promoting Complex (APC/C) and halts progression into anaphase (Maldonado and Kapoor, 2011; Santaguida et al., 2011; Saurin et al., 2011). Once all chromosomes achieve bi-orientation, the SAC needs to be shut down. Inactivation of the SAC, i.e. disruption of the anaphase-inhibitory signal is achieved by removal of key players of the checkpoint, such as Mad1 and Mad2 from the outer kinetochores. This process depends on the microtubule minus-end directed motor Dynein and is known as kinetochore shedding (Basto et al., 2004; Buffin et al., 2005; Griffis et al., 2007; Howell et al., 2001; Mische et al., 2008; Wojcik et al., 2001). Dynein docking at kinetochores is mediated by the ROD-Zwilch-ZW10 (RZZ) complex. The RZZ complex interacts with p50/dynamitin of the dynactin complex and with Dynein intermediate chain, which is regulated by Plk1-dependent phosphorylation (Bader et al., 2011; Whyte et al., 2008). The RZZ complex further contributes to Dynein localisation by interaction with the coiled-coil domain protein Spindly (Chan et al., 2009; Gassmann et al., 2008; Gassmann et al., 2010; Griffis et al., 2007).

Spindly was discovered in a genome-wide RNAi screen for regulators of progression through mitosis in *Drosophila* S2 cells (Griffis et al., 2007). The *Drosophila* Spindly protein has a predicted rod-like structure and contains 780 amino acids forming four coiled-coil domains in the amino-terminal (N-terminal) half and four positively-charged repeats in the carboxy-terminal (C-terminal) half of the protein. In general, the amino acid sequences of Spindly homologues in different species are highly divergent with the exception of a well conserved sequence of twenty-one amino acids called the Spindly box. Spindly localises to the outer kinetochore plate by interacting with the RZZ complex through its C-terminal domain (Barisic et al., 2010; Holland et al., 2015) Mosalaganti et al., 2017; Moudgil et al., 2015). Human Spindly requires farnesylation at cysteine 602 in order to interact with Rod (Holland et al., 2015; Moudgil et al., 2015), while in *C. elegans* Spindly interacts with Zwilch in the absence of hydrophobic interactions (Gama et al., 2017). In addition to the importance of the C-terminal half, other regions of the protein contribute to kinetochore targeting. Human Spindly requires the first coil-coiled domain, the Spindly box and a region between residues 440-564 for kinetochore binding (Barisic et al., 2010).

Once bound to RZZ, Spindly reaches out from the kinetochore to bind Dynein (Barisic et al., 2010; Gassmann et al., 2008; Varma et al., 2013). The first coil-coiled domain of *C. elegans* Spindly is essential for the interaction with the Dynein light-intermediate chain 1 and the Spindly box is important for its interaction with dynactin (Gama et al., 2017). Within the Spindly box, the amino acid residues S256, F258 in human and F199 in *C. elegans* are crucial for the interaction with dynein as their mutation into alanine is sufficient to abrogate the localisation of the motor to kinetochores (Cheerambathur et al., 2013; Gama et al., 2017; Gassmann et al., 2010). In this respect Spindly shares similarities to other Dynein-adaptor proteins such as BicD-2 (Hoogenraad and Akhmanova, 2016; Schlager et al., 2014), that rely on their C-terminal domain to interact with cargoes (i.e. the kinetochore) while the N-terminal portion mediates interaction with the dynein-dynactin motor complex (Griffis et al., 2007). Thus, by linking the RZZ complex with the Dynein motor, Spindly controls cell-cycle progression by acting as a molecular bridge between achievement of bi-orientation and Dynein-dependent checkpoint silencing (Gassmann et al., 2008).

In addition to its role in SAC function, the phenotype of Spindly depleted S2 cells during interphase suggested a role for Spindly in the control of cell shape and the organisation of the cytoskeleton (Griffis et al., 2007). However, a function for Spindly in a tissue or organismal context has thus far remained elusive. Here we analysed Spindly function in *Drosophila* by using loss-of-function and gain-of-function experiments in a tissue-specific fashion. We confirm that a GFP-Spindly construct localises in dynamic fashion during mitosis and that kinetochore localisation requires the C-terminal half of the protein. We find that depletion as well as overexpression of Spindly is embryonic lethal. Despite its requirement for normal localisation in mitosis, deletion of the C-terminal half of the protein only mildly affects Spindly function, while the N-terminal half and Ser234 are essential. In addition to the defects in mitosis due to loss of function, overexpression of Spindly in the germ line results in eggs with altered shape, a mild ventralisation and severely compromised embryogenesis. Finally, we demonstrate that normal levels of Spindly protein are required for migration of the follicular border cells in oogenesis. We conclude that Spindly is required for mitotic functions as well as for movement of post mitotic cells.

## Results

### Requirements for dynamic subcellular localisation of Spindly

Spindly localises dynamically in dividing cultured cells Griffis et al., 2007; Barisic et al., 2010; Gassmann et al., 2008). During prophase and metaphase Spindly associates with the kinetochore and moves poleward in early anaphase. While details of the requirement of Spindly protein domains for its localisation are available for the human homologue (Barisic et al, 2010), this kind of information is still missing for *Drosophila* Spindly. Therefore, in the attempt to perform a domain-function analysis we generated a similar range of GFP-tagged Spindly protein constructs as studied by Barisic et al. (2010) and expressed them in larval neuroblasts using the Gal4/UAS system (Brand and Perrimon, 1993) (**Fig. 1A**). Larval neuroblasts (NBs) are rather large cells, which can be cultured *in vitro,* readily identified and imaged during mitosis at high spatial resolution (Pampalona et al., 2015). For these reasons, larval NBs are an excellent model to study the dynamic localisation of proteins during mitosis. Over-expression of GFP-Spindly in neuroblasts revealed that the protein localises to the kinetochores in a dynamic fashion during mitosis (**Fig 1B**, Suppl.Mat. Movie1). A construct that lacks the C-terminal 380 amino acids did not localise to kinetochores during neuroblast division, indicating that the C-terminal half is important for normal Spindly localisation (**Fig. 1C**; Suppl.Mat. Movie2). All other constructs did localise to the kinetochores (**Fig. 1D,F**; Suppl.Mat. Movie3,4). Another interesting observation was that in 78% of cases (n=40) the protein lacking the N-terminal 228 amino acids localised to cytoplasmic punctae in interphase, which were never observed with any of the other constructs (**Fig. 1E;** Suppl.Mat. Movie5). The speckles do not seem to be polarised in any way. These results indicated that the N-terminal domain is required for the normal distribution of the protein suggesting that the N-terminal deletion causes an abnormal aggregation of the protein in interphase. Furthermore, our experiments confirm that similar to human Spindly, the C-terminal domain is important for binding to the kinetochore.

**Figure 1:**
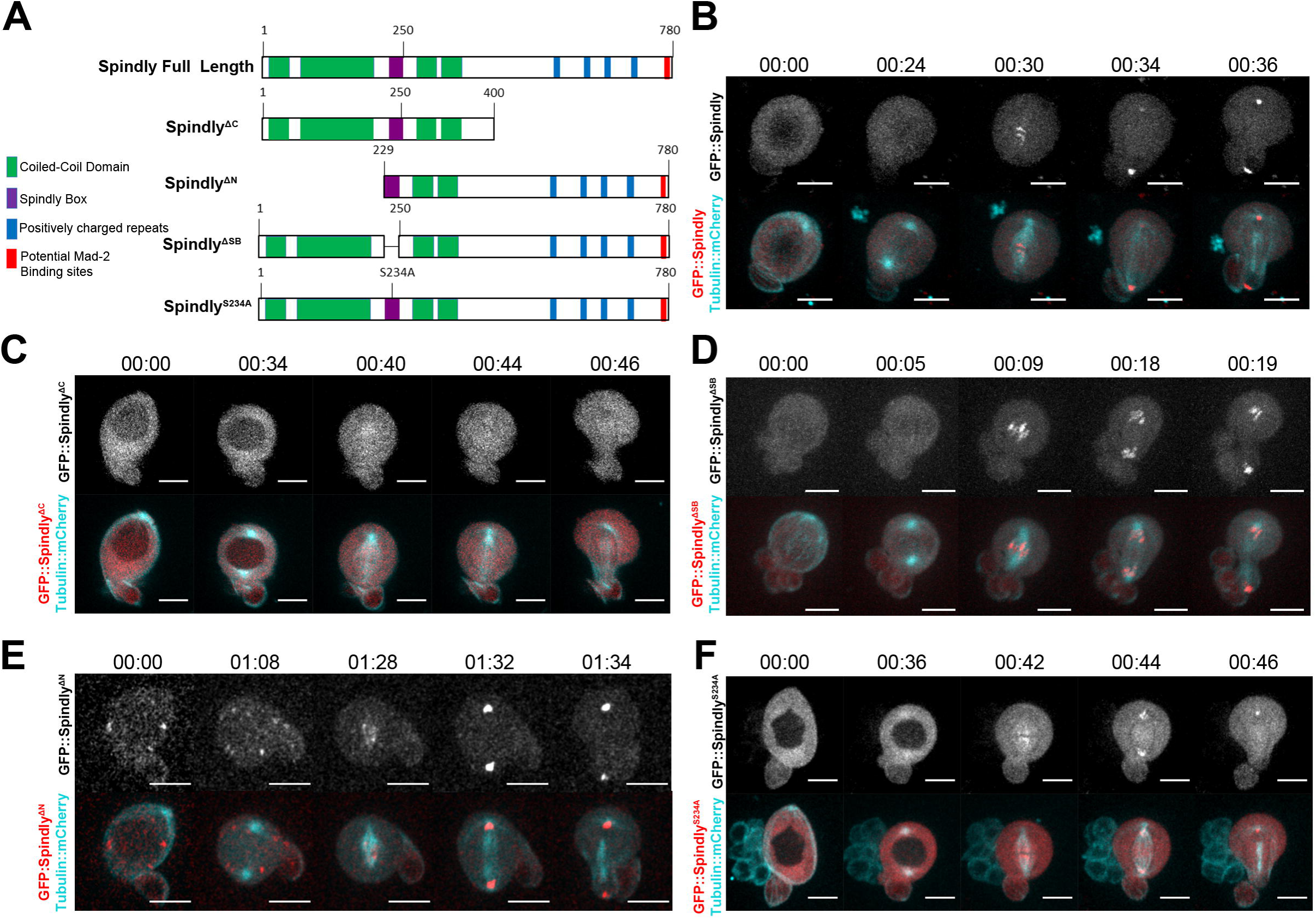
Identification of Spindly protein domain(s) required for dynamic localisation in mitosis. **(A)** Schematic drawing showing the full-length *Drosophila* Spindly protein and mutant constructs generated in this study. Spindly is a predicted coiled-coil domain containing protein (N-terminal half, in green) characterised by a positively-charged C-terminal domain (in blue). The highly conserved repeat motif is depicted (Spindly box, in purple). (**B-F)** Representative examples of dividing neuroblasts overexpressing the indicated *UAS*≫GFP::Spindly variants under the control of the *worniu*>Gal4 driver. **(B)** GFP::Spindly full length gets recruited to kinetochores at early prometaphase (00:26) and can be seen moving from KT to poles at anaphase (00:32) In all cases (n=21) the protein localised this way. **(C)** GFP::Spindly^Δct^ never localised at kinetochores throughout mitosis (n=43). Mutant constructs deleted of the Spindly box (n=21) **(D)**, N-terminal half (n=40) **(E)** or a point mutant S234A (n=33) **(F)** showed localisation dynamics similar to the full length protein. Note the speckled pattern of GFP::Spindly^ΔNt^ in interphase, which was observed in 31/40 cases. Time stamp: hh:min. Scale bars: 10 µm.

### RNAi knock-down of Spindly in *Drosophila* development

Since Spindly knock-down by RNAi worked efficiently in *Drosophila* S2 cells (Griffis et al., 2007), we decided to use RNAi resources and selected shRNAs directed against *spindly* mRNA that can be expressed under temporal and spatial control by the Gal4/UAS system (Brand and Perrimon, 1993; Ni et al., 2009; Ni et al., 2008).

A maternal Gal4 driver (alpha-*tubulin67*>*Gal4*) was used to express UAS-controlled shRNA targeting *spindly* mRNA in oocytes and embryos (called *spindly*[mat67>RNAi] hereafter). The efficacy of the transgenic RNAi knockdown was assayed by immunoblotting of embryo lysates with an affinity-purified Spindly antibody, and a marked reduction in the levels of Spindly protein was observed at all temperatures tested (**Fig. 2A**). This result indicated that the Gal4-driver used caused RNAi-dependent depletion of the protein encoded by the targeted mRNA at all temperatures. We did not observe a stringent correlation between the efficacy of RNAi using this driver at different temperatures, probably due to the severe abnormalities of the embryos from very early on.

**Figure 2:**
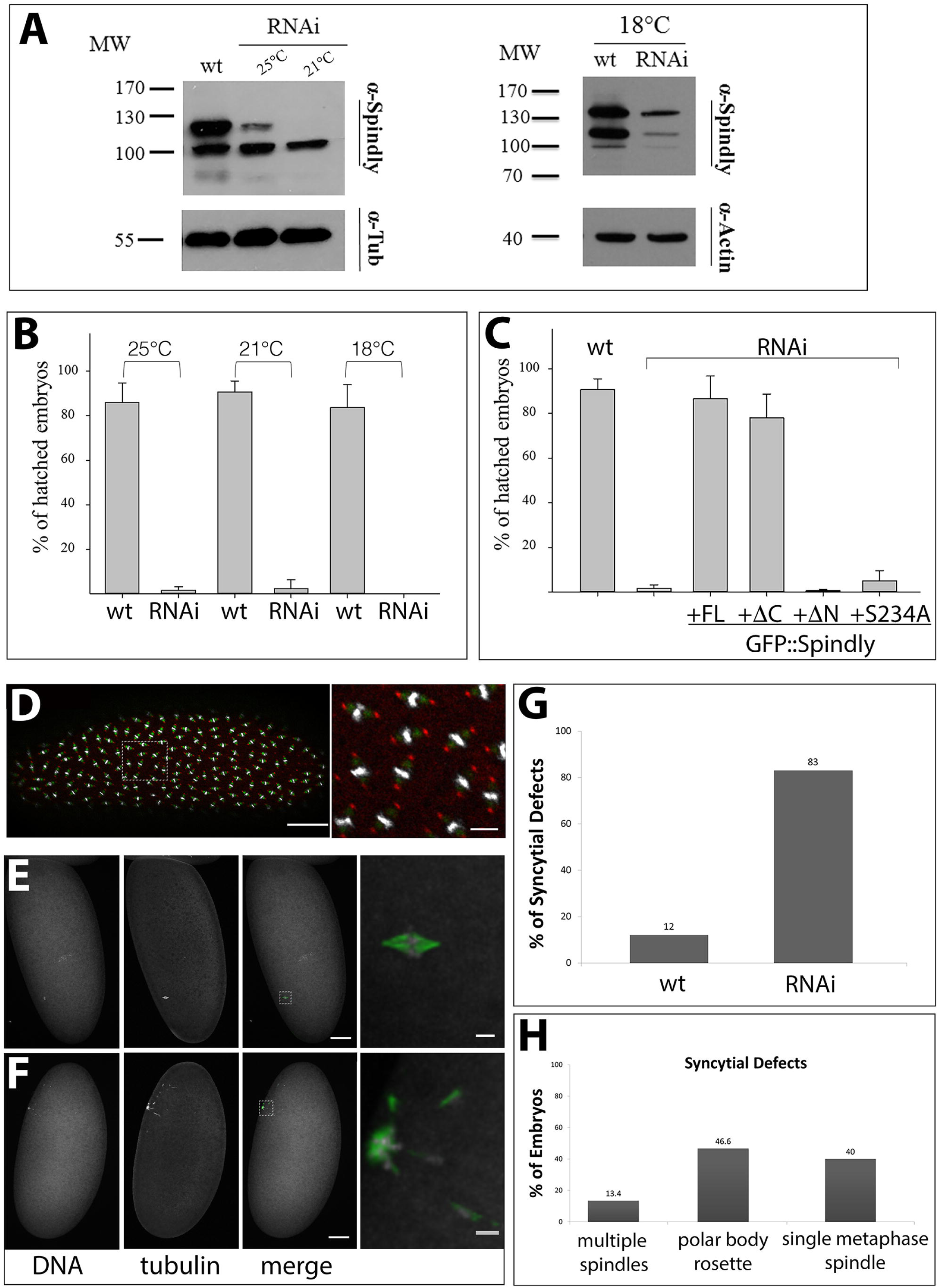
*mat-tub*≫*Gal4* driven Spindly knock-down. **(A)** Western blot analyses of protein lysates from 0-3 hours wild-type (wt) and *spindly mat67*≫RNAi embryos (RNAi) using anti-Spindly antibody. Embryos were cultured at different temperatures from 25°C to 18°C as indication (antitubulin or anti-actin were used as loading control; Mw - Molecular weight (in kDa)). **(B)** Larval hatching rates of *spindly*[*mat67*≫RNAi] embryos. (n=100) **(C)** Larval hatching rates after expression of GFP-Spindly in wild type (wt; *w*^1118^), *spindly*[*mat67*≫RNAi] (RNAi) and *spindly*[*mat67*≫RNAi] embryos expressing either GFP::Spindly full-length (+FL), GFP::Spindly^ΔCt^ (+ΔCt),GFP::Spindly^ΔNt^(+ΔNt) or GFP::Spindly^S234A^ (+S234A) (n=100). **(D)** Mitotic domains in wild-type syncytial blastoderm embryo. Nuclei are stained with DAPI (grey), tubulin (green) and anti-Centrosomin (red). Scale bars represent 50 µm and 10 µm. **(E,F)** Syncytial division defects in *spindly mat67*≫RNAi embryos. The images are maximum-intensity projections of 10 μm z-stack (at spacing of 1 μm). (**E**) Embryos often contained a single metaphase-arrested nucleus. (**F**) In some cases a polar body rosette was visible (DAPI (grey) and anti-tubulin (green); scale bar: 50 μm; details: 5 μm. (**G**) Penetrance of syncytial division defects in *spindly mat67*≫RNAi embryos (n=100). **(H)** Expressivity of syncytial cleavage defects in *spindly mat67*≫RNAi embryos (n=30).

*spindly*[mat67>RNAi] had severe effects on embryogenesis causing a remarkable embryonic lethality (**Fig. 2B**). To exclude that the effects of the knock-down were caused by off-target effects we generated an RNAi-resistant construct encoding an N-terminally GFP-tagged full-length version of Spindly in which the shRNA target site was mutated without changing the primary polypeptide sequence. The over-expression of the RNAi-resistant GFP-tagged Spindly transgene fully restored the viability of *spindly*[mat67>RNAi] embryos to wild-type levels (**Fig. 2C**). The requirements for different protein domains was tested by expressing the RNAi-resistant GFP constructs introduced above (**Fig. 1A**) in *spindly* [mat67>RNAi] embryos. We found unexpectedly that the C-terminal half was largely dispensable for the rescue of *spindly*[mat67>RNAi] embryos, while the N-terminal half and the amino acid residue Ser234 within the Spindly box were required for the function of the protein (**Fig. 2C**). Thus knock-down of *spindly* mRNA using transgenic RNAi to maternally express shRNA provides a suitable model to investigate Spindly function in *Drosophila*. Our results suggest that kinetochore localisation is not critical for Spindly function in embryos.

### Requirement of Spindly in syncytial cleavage division

After fertilisation, fly embryos undergo rapid nuclear mitotic division cycles in a common cytoplasm. The well-established function of Spindly for mitosis suggested that *spindly*[mat67>RNAi] should affect these syncytial divisions potentially explaining the embryonic lethality observed upon its depletion (**Fig. 2B**). Indeed, immunostaining of mitotic spindles in early embryos revealed that, in contrast to controls, 83% (n=100) of the *spindly*[mat67>RNAi] embryos showed abnormal syncytial cleavage divisions (**Fig. 2D-H**). The majority of embryos arrested during metaphase of the first 1–3 mitotic divisions (**Fig. 2E,F,H**). In 40% of the embryos (n=30), only a single metaphase-arrested mitotic spindle is present within the egg (**Fig. 2E**) whereas only in 13% of the cases a variable number of 2 to 4 mitotic spindles were observed (**Fig 2H**). We also detected characteristic polar body rosettes (Page and Orr-Weaver, 1997) in 46.6% of embryos suggesting that meiosis is able to resume and complete in *spindly*[mat67>RNAi] embryos (**Fig. 2F**). We conclude that Spindly is essential for mitotic cycles from the first syncytial cleavage division on.

To analyse the requirement of Spindly during later mitotic cleavage divisions, we drove *spindly* RNAi using the *nanos* (*nos*)-Gal4 driver (hereafter *spindly*[*nos*>RNAi]), which drives germ-line expression in the germarium. *spindly*[*nos*>RNAi] was effective in knocking down Spindly, but in contrast to [mat67>RNAi] also exhibited a remarkable temperature sensitivity (**Fig. 3AC**). At 21° to 25°C *spindly*[*nos*>RNAi] efficiently depleted Spindly protein in early embryos, being most efficient at 23° to 25°C (**Fig. 3A**). Consistent with the depletion of Spindly protein, *spindly*[*nos*>RNAi] embryos showed also temperature-dependent rates of survival. Compared to the wild-type only 16% (n=100) of embryos were able to hatch at 23°C, while at 21°C about half of the *spindly*[*nos*>RNAi] embryos survived to larval stages (n=100) (**Fig. 3B**). Immunostaining of *spindly*[*nos*>RNAi] embryos raised at either 23°C or 21°C was performed to further analyse the cause of the embryonic lethality. DAPI staining of *spindly*[*nos*>RNAi] embryos and quantification of syncytial cycle defects were consistent with a requirement for Spindly in normal embryonic development (**Fig. 3C**).

**Figure 3:**
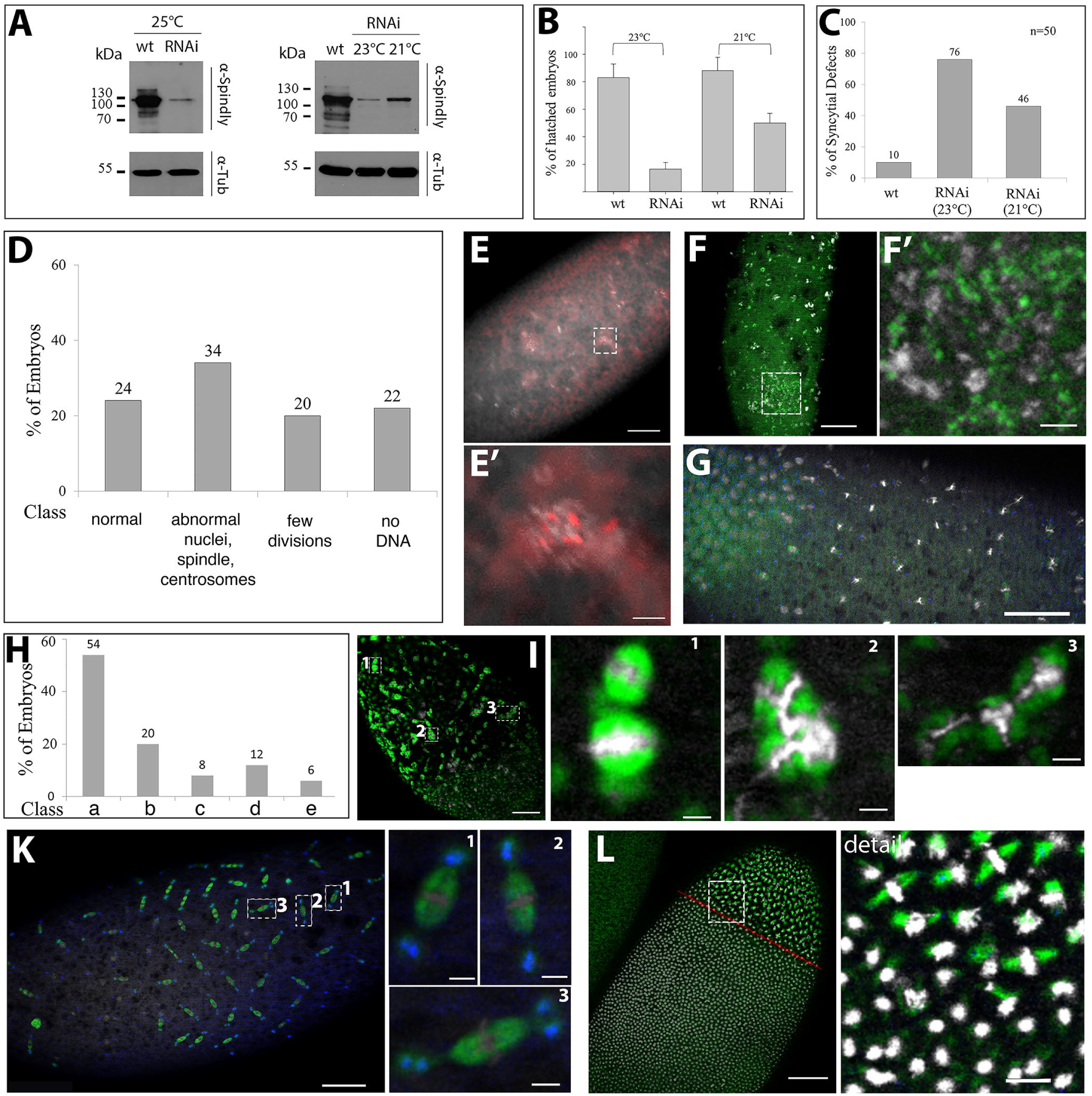
*nos*≫Gal4 driven Spindly knock-down. **(A)** Immunoblot analysis of protein lysates of 0-3 hours old *w*^1118^ embryos (wt) or *spindly*[*nos*>RNAi] (RNAi) embryos at different temperatures (loading control - α-Tubulin (Tub); molecular weight standards in kDa). **(B)** Viability of *spindly*[*nos*>RNAi] scored by larval hatching rate (n=100). **(C)** Penetrance of syncytial defects of *spindly*[*nos*>RNAi] embryos inferred from DNA staining of fixed embryos. **(D)** Expressivity of syncytial cleavage defects of *spindly*[*nos*>RNAi] embryos at 23°C. Occurrence of phenotypic classes of *spindly*[*nos*>RNAi] embryos (n=50). **(E)** Syncytial stages with patches of disorganised chromatin occasionally associated with multiple centrosomes; (**E’)** higher magnification of boxed area in (**E**); (DAPI (gray), centrosomes (red); scale bars: 50 µm (E) and 10 µm (E’)). (**F**) Condensed chromatin associated with patches of tubulin (detail indicated by yellow box; DAPI (grey), anti-tubulin (green), scale bars represent 50 µm (F) and 10 µm (F’)). (**G**) Asynchrony of mitotic divisions indicated by area of mitotic divisions and areas enriched in interphase nuclei (scale bar represents 50 µm). **(H)** Expressivity of syncytial cleavage defects of *spindly*[*nos*>RNAi] embryos. *spindly*[*nos*>RNAi] embryos raised at 21°C displayed a range of mitotic defects (n=50): (b) uneven spacing of the mitotic domains and asynchrony of nuclear divisions. (c) abnormal mitotic figures. (d) centrosome abnormalities (detachment, monopolar spindles, centrosome clustering). (e) dead embryos a later stages. (**I**) Abnormal, irregular spacing between adjacent mitotic domains. Details are shown from the dashed yellow boxes in merged panel: (**1**) mitotic spindles collapse and fuse; (**2**) abnormal tubulin-rich structures associated with chromatin; (**3**) tri-polar spindle. (Scale bars represent 50 μm and 5 μm (for details)). **(K)** Defects in cohesion between centrosomes and spindle poles. Details from merged panel marked with a dashed yellow box: (**1**) a pair of centrosomes associated to a single spindle pole. Detail 2 and 3: centrosomes detach from one (**2**) or both poles (**3**); red arrows point to detaching centrosomes. Scale bar represents 50 μm and 5 μm (for details). **(L)** embryo with domains of distinct mitotic rate (red line marks boundary between a region of interphase nuclei and a region with mitotic divisions; scale bars represent 50 μm and 10 μm (detail).

Fluorescence microscopy was used to assess any mitotic defects in *spindly*[*nos*>RNAi] embryos by analysing centrosome positioning and mitotic spindle morphology. At 23°C, the majority of *spindly*[*nos*>RNAi] embryos had arrested before the end of syncytial divisions (76% n=50; **Fig. 3D**). 22% of all embryos did not show any DAPI staining while 20% arrested development after a few rounds of syncytial mitotic cycles. Of this latter category, 10% of the embryos were characterised by patches of disorganised chromatin associated with multiple centrosomes (**Fig. 3E,F**). The normal organisation of the mitotic spindle was compromised and condensed chromatin was found associated with patches of tubulin (12%, n=50; **Fig. 3F**). Unlike wild-type embryos, the nuclei of *spindly*[*nos*>RNAi] embryos were unequally distributed in the cortical cytoplasm of the embryos and the regular spacing of the mitotic domains was lost (**Fig. 3G**). In summary, at the highest temperature tested, *spindly*[*nos*>RNAi] resulted in severe disorganisation of centrosome distribution and chromatin structure in early syncytial stages.

At 21°C only just under 50% of *spindly*[*nos*>RNAi] embryos failed to hatch and exhibited defects in the synchrony of syncytial divisions (see below) and various spindle defects (**Fig. 3H**). The most severe spindle defects might result from abnormal spacing of the mitotic domains in the cortex of the syncytial embryo. Often mitotic spindles appeared to be stacked together, fusing into highly abnormal microtubule-based structures (**Fig. 3I).** About 20% of the abnormal embryos exhibited various centrosome defects, including centrosomes detaching from one or both spindle poles or a pair of centrosomes associated with one or both poles (**Fig. 3K**). The phenotypes suggest that centrosomes might undergo abnormal replication or they detach and move away from the spindle pole or that free centrosomes occur by nuclear displacement away from the cortex and leaving centrosome behind. The results suggest that lower levels of *spindly*[*nos*>RNAi] mediated knockdown produced a threshold of Spindly activity that uncovers its role in spindle assembly and maintenance, which may go beyond its functions as a component of SAC.

In the early embryo, the observed fusion of mitotic spindles suggests a function of Spindly in the separation of mitotic domains during cortical syncytial divisions. A characteristic feature of *spindly*[*nos*>RNAi] syncytial embryos was a defect in synchrony of syncytial divisions. *spindly*[*nos*>RNAi] contained areas with different rates of mitotic activity, i.e. part of the embryo contained interphase nuclei, while in adjacent areas nuclei were still in mitotic phases (**Fig. 3G,L**). This asynchrony in mitotic divisions and uneven spacing of nuclei was observed at all temperatures examined and suggests that limitations of Spindly protein levels blocks the wave-like progression of mitotic divisions in the syncytial embryo.

### Spindly overexpression cause defects in egg shape and embryogenesis

The loss-of-function studies reported here demonstrated that maternally supplied Spindly is essential for sustaining the early mitotic cell cycles in the syncytial embryo. Importantly, the phenotypes were specific for Spindly RNAi knockdown, as a full rescue of the defects was observed by expression of an RNAi-resistant Spindly protein. In the course of these experiments we noted that expression of GFP-Spindly using maternal *tubulin67c>Gal4,* in a wild-type background caused defects in embryogenesis. In *mat67>>GFP-Spindly* embryos, levels of Spindly-GFP were higher than endogenous protein levels in lysates from 0-3 hours-old embryos at 25°C (**Fig. 4A**).

**Figure 4:**
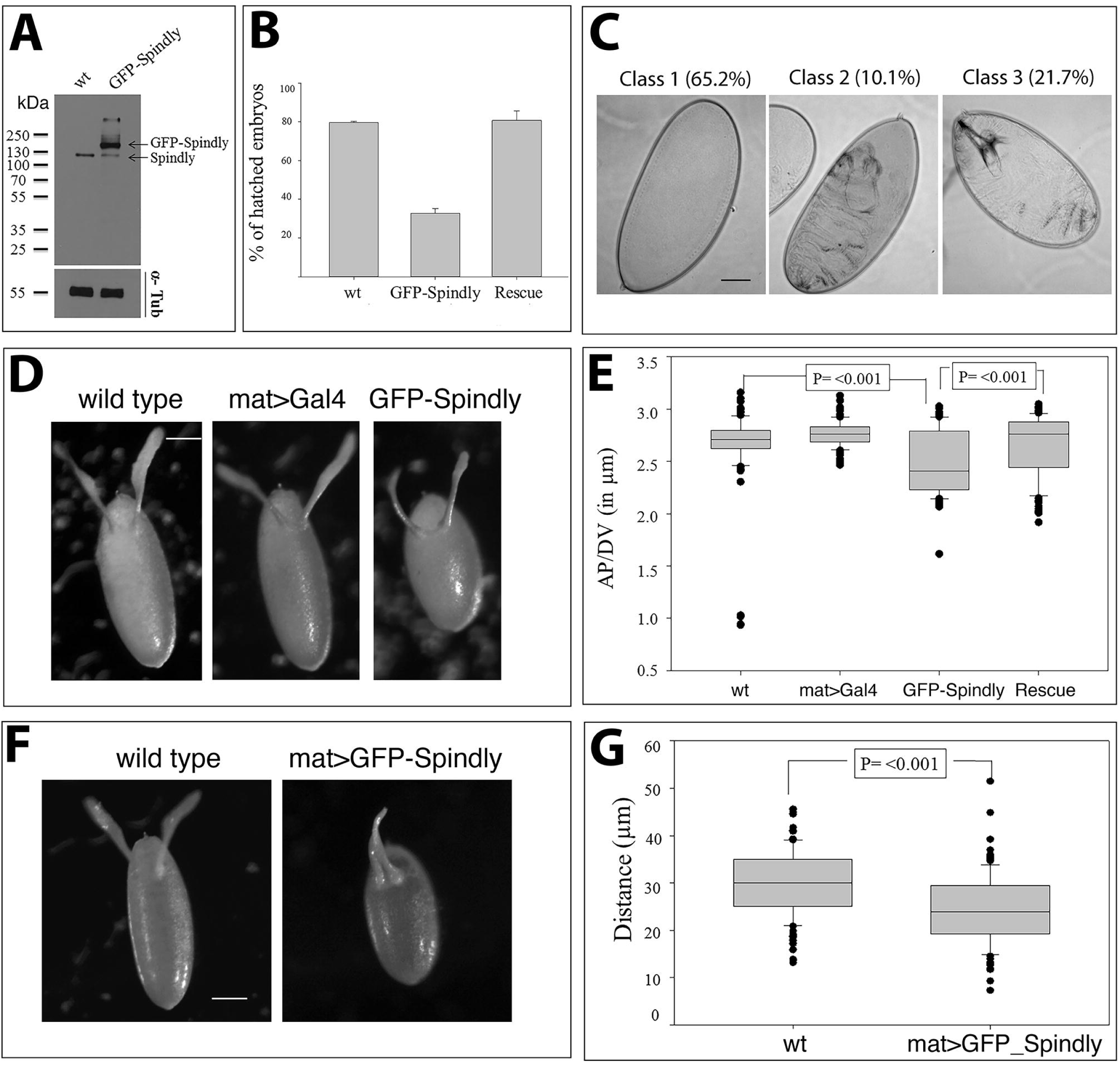
Over-expression of Spindly affects oogenesis and embryogenesis. (**A**) Immunoblot analysis of Spindly in lysates from wild-type (*wt*) and *nos≫Gal4*:*UAS≫GFP-Spindly* (GFP-Spindly) embryos. Endogenous Spindly and GFP-Spindly bands are indicated (α-tubulin - loading control; molecular weight markers in kDa). (**B**) Survival rate of embryos upon Spindly overexpression scored by larval hatching (n=100). Viability was restored by knock-down of endogenous Spindly with UAS≫Spindly[RNAi] (Rescue). (**C**) Larval cuticle defects in *mat67≫Gal4*:*UAS≫GFP-Spindly.* The majority of embryos died before cuticle deposition (65.2%, class 1). The remaining 31.8% of embryos exhibited dorsal holes (class 2) or segmentation defects (class 3) (n=138). (**D**) Eggs laid by *mat67≫Gal4*::*UAS≫GFP-Spindly* females exhibited abnormal rounded shape (48%, n=100). Scale bar represents 50 μm. (**E**) Quantification of egg size by calculating the ratio between the length (A/P axis) and the width (D/V axis) for a total of 100 eggs (Mann-Whitney Test (p=<0.001). This phenotype is suppressed by UAS≫Spindly[RNAi] knock-down of endogenous Spindly (Rescue). (**F**) Moderate ventralisation defect in chorions of *mat67≫Gal4*::*UAS≫GFP-Spindly* eggs. The dorsal appendages were shorter and closer apposed to each other along the dorsal midline compared to controls. Scale bar represents 50 μm. (**G**) Quantification of the distance between dorsal appendages shown in (**F**) (Mann-Whitney Test p=<0.001; n=100).

The majority (70%, n=100) of *mat67>>GFP*-*Spindly* embryos did not hatch, but co-expression of Spindly[shRNA] was able to fully rescue this effect indicating that this phenotype was caused by the altered level of Spindly protein rather than by an abnormal effect through expression of the GFP-Spindly fusion protein (**Fig. 4B**). The embryonic lethality of *mat67>>GFP-Spindly* resulted in three classes of larval cuticle phenotypes (**Fig. 4C**). Within the largest class (65%) embryos did not form any cuticle suggesting that the embryos died early, while the remaining embryos exhibited cuticle phenotypes suggesting a range of morphogenetic defects (**Fig. 4C**). In addition to embryonic defects, normal egg shape was impaired by *mat67>>GFP-Spindly* expression resulting in eggs that were shorter along their anterior-posterior axis (**Fig. 4D,E**). The morphology of the dorsal-anterior appendages of the chorion was also affected in a way that suggested a mild ventralisation phenotype as the distance between the attachments of the appendages was significantly shorter in *mat67>>GFP-Spindly* eggs (**Fig. 4F,G**). These data indicate maternal over-expression of Spindly protein affects egg formation and patterning in oogenesis and embryonic development suggesting that spatio-temporal regulation of *spindly* expression and/or protein levels are crucial for survival.

### Requirement of Spindly in ovarian border cell migration

Knockdown of Spindly drastically alters S2 cell morphology in interphase: Spindly-RNAi cells have malformed actin lamellae and abnormal microtubules arranged in long bundles that protrude from the cell body (Griffis et al., 2007). Furthermore, overexpression of a GFP-tagged version of the protein revealed its accumulation at the (+) ends of microtubules along with other microtubule-plus-end binding factors such as EB-1 (Griffis et al., 2007). We therefore addressed whether Spindly has a role in regulating the motile behaviour of cells.

The border cells (BCs) in the *Drosophila* ovary provide an excellent model system to study genetic requirements of collective cell migration (Montell, 2003). The BCs form a cluster of 6-10 post-mitotic cells at the anterior domain of somatic follicular epithelium of the egg chamber around stage 8 of egg chamber development (**Fig. 5A**). Subsequently, BCs undergo an epithelial-mesenchymal transition and migrate posteriorly as a coherent cluster reaching the nurse cell-oocyte boundary by stage 10 of egg-chamber development (Montell, 2003); **Fig. 5A**). Spindly expression was knocked down by driving shRNAi with the BC-specific *slow border cells* (*slbo*) Gal4 driver (*slbo>Gal4*) (Rorth, 1994). In ovaries from *slbo*>>Spindly[RNAi] females, 50% (n=46) of the BC clusters had completed migration at mid-stage 9 of oogenesis (**Fig. 5B,C**). Conversely, in wild-type egg chambers the majority of BC clusters had migrated half way through the egg chamber at late stage 9 (69%, n=42), while a small percentage (14.3%) had already completed their migration at this stage of egg-chamber development (**Fig. 5B,C**). Normalised migration (MI) and completion (CI) indices were calculated to compare BC cluster migratory phenotypes upon Spindly-RNAi according to (Assaker et al., 2010). Values of both indices suggest that down-regulation of Spindly protein level confers to BC a migratory advantage (**Fig. 5B**). Thus, we concluded that knockdown of Spindly caused the BC cluster to migrate either earlier or faster along the anterior-posterior axis towards the oocyte suggesting that Spindly negatively regulates border cell migration.

**Figure 5:**
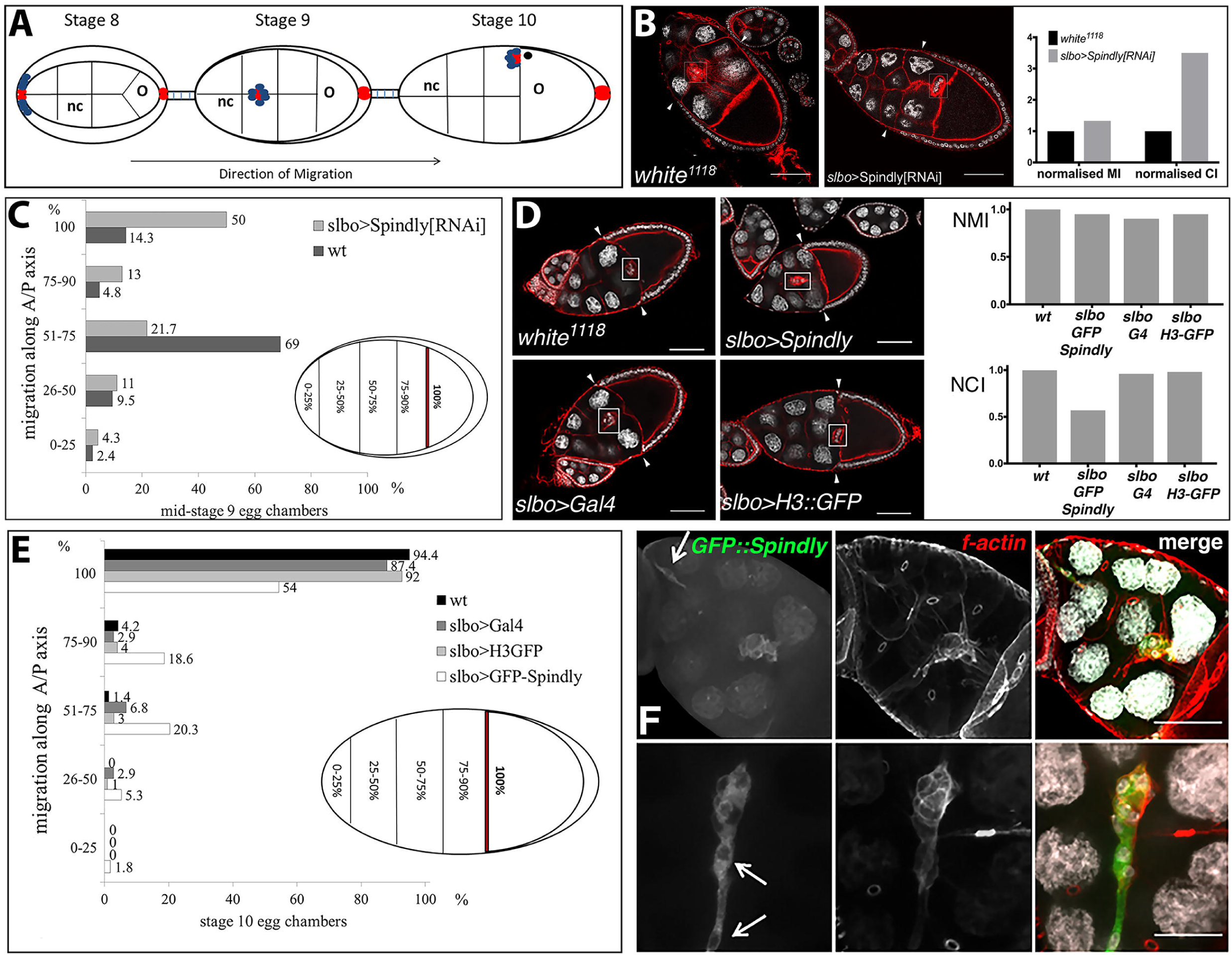
Spindly in border cell migration. (**A**) Schematic drawing of border cell migration and outer follicle cells rearrangement. Egg chambers are depicted at stage 8-10 of oogenesis, where each egg chamber consists of 16 germ line cells (15 nurse cells (nc) and one oocyte (o)) and are surrounded by a layer of somatic follicle cells. At stage 8, polar cells (in red) at the anterior of the follicular epithelium recruit neighbouring cells (in blue) to form the migrating border cell cluster. At stage 9, the cluster delaminates from the epithelium and starts migrating posteriorly towards the oocyte and arriving at the nurse-cell-oocyte boundary at stage 10. (**B**) Immunostaining of stage 9 egg chambers from wild-type (*w*^1118^) and *slbo>Gal4;UAS>SpindlyRNAi* females (Nuclei: DAPI (grey), f-actin (red); Scale bars: 50 μm; MI-migration index; CI-completion index). (**C**) Quantification of the phenotype described in (**B**). Premature border cell migration was observed in 50% of stage 9 egg-chambers (n= 42). (**D**) Confocal optical sections of egg-chambers at stage 10 of oogenesis of the indicated genotypes. Migration of border cells was delayed by GFP-Spindly overexpression (DAPI (grey); f-actin (red); scale bars - 50 μm; NMI-normalised migration index; NCI-normalised completion index). (**E**) Quantification of the phenotype described in (**D**). 46% of the clusters overexpressing Spindly did not complete migration by stage 10 of egg-chamber development (n= 113). (**F**) Morphology of border cell clusters in egg chambers of *slbo>Gal4; UAS>GFP-Spindly* females. Confocal optical section of stage 10 egg-chambers (maximum-intensity projection of 10 μm z-stack at a spacing of 1 μm). Loss of border cell integrity is marked by white arrows. The phenotype was observed in 9.7%; single cells detached from the main cluster and migration was delayed (**F**, upper row panels). In 1.8% of cases dissociation of BC clusters was observed (**F**, lower row panels) (n= 113) (DAPI (grey), f-actin (red), GFP (green); scale bars represent 20 μm).

The sensitivity of BCs to Spindly levels prompted us to test the effect of Spindly overexpression on their migration. Specifically, we asked whether overexpression of Spindly could change BC migration at stage 10 of egg chamber development. Indeed, when analysing migration of the border cells, we found that in 46% (n= 113) of BC clusters their migration towards the oocyte was delayed at stage 10 of egg-chamber development. While *slbo*>Gal4 females on their own also exhibited a mild incomplete migration phenotype at stage 10 (12.6%; n= 103), the penetrance of the phenotype was significantly lower compared to *slbo>>GFP-Spindly* overexpression (Fig. **5D,E**). As an additional control, overexpression of GFP-tagged histone H3 did not alter border cell migration (n=100), supporting the result that the increased Spindly expression causes incomplete border cell migration (Fig. **5E**).

Overexpression of Spindly in BCs also affected their morphology in a way suggesting abnormal adhesive properties and cytoskeletal architecture. Many of the BC clusters lost their integrity and the cell-cell contacts between BCs were compromised to the extent that individual cells detached from the BC cluster and thus were apparently migrating as individual cells (9.7%, n=113) (**Fig. 5F**). These results suggest that excess Spindly negatively affects the migration and the integrity of the border cell cluster during migration. We conclude that correct Spindly levels are required in post-mitotic follicle BC to control or modulate their migration possibly by controlling integrity and adhesion of the cluster.

## Discussion

Since its discovery in 2007, Spindly has been well characterised as a crucial component of the mitotic checkpoint by coordinating chromosome alignment to SAC inactivation (Griffis et al., 2007) (Chan et al., 2009) (Gassmann et al., 2010) (Barisic et al., 2010). With the exception of two studies in *C. elegans* Spindly has been investigated using tissue culture cell lines(Gassmann et al., 2008). Here we used the Gal4-UAS system to characterise the mitotic and post-mitotic activities of *Drosophila* Spindly in a variety of cellular and developmental processes. We found that loss of Spindly compromises embryonic development by altering mitotic spindle morphology and synchrony of the syncytial mitotic cleavages. Likewise, increasing the expression level of Spindly above the physiological threshold results in embryonic lethality, altered morphology and moderate ventralisation of the eggs. Finally, our study uncovers a novel and unexpected function for Spindly in post mitotic cells, regulating the migration of the border cell cluster.

### Spindly function in early *Drosophila* embryo development

The temperature sensitivity of the Gal4/UAS system provided a means to generate embryos with different maternally-derived levels of Spindly protein. Consistent with its central role in the mitotic checkpoint, *spindly*[RNAi] embryos exhibit abnormal mitosis, with phenotypes ranging from mild to severe as would be expected from a series of hypomorphic to amorphic mutant alleles. Analysis of nuclear proliferation in *spindly*[RNAi] embryos revealed that all embryos showed abnormal mitotic progression. Severe knock-down of Spindly blocked embryogenesis at the first mitotic divisions. The arrest took place either at the first or second nuclear division cycle and suggests an important role for Spindly in the control of the transition from a mature oocyte into cleavage stages of embryonic development. A possible cause of this phenotype could be a failure in the fusion of the female and male pronuclei. Given the severe disruption of the microtubule cytoskeleton observed in early Spindly[RNAi] embryos, Spindly might contribute to the assembly of the microtubule network that is nucleated by the male aster and necessary to guide the female pro-nucleus movement towards the male pronucleus (Loppin et al., 2015).

Embryos with lower knock-down of Spindly displayed an uneven spacing and asynchronous division of nuclei, mitotic spindle defects, as well as detached and supernumerary centrosomes. Similar asynchrony defects are observed in *Drosophila* embryos injected with Dynein inhibitors (Sharp et al., 2000b). These observations therefore point to a more general inhibition of dynein activity and support the idea of Spindly being a more general regulator of this motor protein. Moreover, it is well known that the maintenance of the spacing and architecture of the spindle machinery is achieved by the coordinated and complementary activity of different spindle-associated Dynein and Kinesin motor proteins (Sharp et al., 2000a) (Wang et al., 2014). Loss of Spindly function might alter the balance between the opposing forces that govern spindle stability, resulting in the altered spindle morphology and chromosome segregation defects described in this work. Finally, the crucial role of Dynein in maintaining a tight association between the centrosomes and the spindle poles has been extensively described in early *Drosophila* embryos as well as in S2 cells (Robinson et al., 1999) (Morales-Mulia and Scholey, 2005). It is therefore appealing to hypothesise that the phenotypes described in this report reflect a role for Spindly in coordinating kinetochore and non-kinetochore functions of Dynein during mitosis. Our observations corroborate the idea that Spindly must interact with and assist Dynein to ensure proper execution of mitosis and embryonic development.

Spindly forms a ternary complex with Dynein and the Dynactin complex in order to fulfil its functions (Griffis et al., 2007) (Gassmann et al., 2008) (Gassmann et al., 2010) (Gama et al., 2017). The first coiled-coiled domain in the N-terminal portion of the protein has been shown to be responsible for the interaction with Dynein LIC-1 (Gama et al., 2017). Moreover, highly conserved residues within the Spindly box, residues S256A, F258A in human and F199A in *C. elegans*, are essential for the interaction with the Dynactin complex and for the binding of dynein at kinetochores (Cheerambathur et al., 2013; Gama et al., 2017; Gassmann et al., 2010). Our observation that expression of Spindly^ΔNt^ and SpindlyS^234A^ in a *spindly*[RNAi] background did not rescue embryo viability suggests a similar importance for these domains for the interaction with Dynein. Although the interaction between *Drosophila* Spindly and Dynein has yet to be demonstrated, our results support the idea that knock-down of Spindly reflects a loss of Dynein function in establishing and maintaining stable kinetochore-microtubule interactions (Gassmann et al., 2008).

We confirmed earlier studies on human Spindly by showing that the C-terminal domain of *Drosophila* Spindly is required for its localisation at kinetochores in mitosis (Barisic et al., 2010; this study). To our surprise the overexpression of a Spindly^ΔCt^ construct, which is unable to localise to kinetochores did rescue embryo viability. If the activity of Spindly was exclusively kinetochore-dependent, overexpression of Spindly^ΔCt^ would be expected to further enhance the phenotype of Spindly[RNAi]. We therefore propose that Spindly likely has additional functions including kinetochore-independent functions *in vivo*.

Spindly overexpression led to embryonic lethality and revealed defects in egg morphology, shape, size and a moderate ventralisation phenotype. The establishment of dorsal/ventral polarity occurs during mid-oogenesis and ultimately relies upon a dorsalising signal generated by the *gurken* gene (Schupbach, 1987) (Forlani et al., 1993). The mild dorsal/ventral patterning defects of eggs over-expressing Spindly could be interpreted as a consequence of a displacement of the oocyte nucleus and consequently of the fate determinant Gurken away from the usual anterior-dorsal position. Nuclear migration during oogenesis and anchoring of the oocyte nucleus to the anterior cortex are both events that rely on the motor protein Dynein and the activity of its co-factors (Duncan and Warrior, 2002; Januschke et al., 2002; Lei and Warrior, 2000; Sitaram et al., 2014; Swan et al., 1999; Swan and Suter, 1996). The Spindly overexpression phenotype is therefore also consistent with the idea of Spindly acting as a Dynein co-factor in multiple developmental processes, including egg patterning.

### A role of Spindly in cell migration

In the field of mitosis research, Spindly is seen as an important coordinator of the activity of Dynein by regulating the subcellular localisation, directionality and processivity of the motor (McKenney et al., 2014) (Schlager et al., 2014). In addition to its role as Dynein cofactor, previous work suggested a role for Spindly in cytoskeleton remodelling (Griffis et al., 2007). *Drosophila* Spindly localises at the (+) ends of microtubules in interphase S2 cells (Griffis et al., 2007) and its downregulation results in a peculiar phenotype in interphase, with cells displaying severe defects in the lamellae network as well as in the overall organisation of the microtubule cytoskeleton. The loss- and gain-of-function experiments performed on follicular border cells have opened an ideal background to test the role of Spindly in the regulation of cell motility. We found that downregulation of Spindly positively regulates the migration of border cells towards the oocyte. Conversely overexpression of GFP-tagged Spindly resulted in an incomplete migration phenotype and abnormal cluster morphology. The Dynein light chains, Lis-1 and Nud-E have been implicated in border cell migration; however in contrast to Spindly-RNAi, knock-down of these proteins delayed border cell migration (Yang et al., 2012). In addition, Dynein chains are required for the normal organisation of the cluster, proper localisation of adhesion molecules and epithelial cell polarity (Yang et al., 2012) (Horne-Badovinac and Bilder, 2008). These data suggest that altering the activity of dynein might cause defects in border cell cluster morphology. In the context of the gain-of-function experiments we can hypothesise that overexpression of Spindly might have similar consequences to the overexpression of Dynamitin (Quintyne et al., 1999) (Schroer, 2000), resulting in the inhibition of dynein activity and altered distribution of polarity markers and/or adhesion molecules. This interpretation would nicely fit with the mild-ventralisation of the egg upon Spindly overexpression, a phenotype that could arise from reduced Dynein activity and therefore the detachment of the oocyte nucleus from the anterior-dorsal cortex.

### Concluding remarks

In conclusion, we find that Spindly is required for distinct cellular functions which we can expose by changing the levels of Spindly protein using modulatory RNAi or overexpression scenarios. The different loss and gain-of-function phenotypes reported here can be explained by postulating a “threshold model” for Spindly activity. For a multifunctional protein as Spindly, a certain biological function would be performed only when the levels of protein expression hits a certain threshold. This implies that Spindly could have different local activity thresholds for distinct mitotic and non-mitotic functions. Therefore, certain levels of knockdown or overexpression would produce a certain category of defects. This model implicates that the expression levels of Spindly must be under tight control for the protein to properly fulfil its multiple activities.

## Materials and methods

### Drosophila strains and culture

Flies were kept on standard medium and grown accordingly to standard laboratory procedures. *white*^*1118*^ serves as wild-type control strain (*Bloomington Stock Centre*). *To express genes ectopically* and for the RNAi experiments, the Gal4-UAS system was used (Brand and Perrimon, 1993). The Gal4 lines used were: *nanos*>Gal4 VP16 (Ulrike Gaul), *maternal tubulin 67c*>Gal4, *slbo*>Gal4 (Denise Montell). The UAS lines used were p*UASp*>GFP-Spindly constructs (this work), UAS>Spindly RNAi (34933 *Bloomington Stock Centre*), *UAS>H3-GFP*, worniu-GAL4 (Albertson et al., 2004), and UAS-mCherry::tubulin (Rusan and Peifer, 2007). Transgenic flies harbouring the p*UASp*>GFP-Spindly construct were generated by PhiC31– mediated sequence-directed transformation using an attP site at the genomic location 68A4 on the third chromosome.

### Embryo collection

Embryos were collected on yeasted apple juice agar plates. To determine hatching rates, embryos from an overnight collection were counted and aged at 25°C, 23°C, 21°C, 18°C for 48 hours. The number of hatched embryos was determined by counting the number of intact, unhatched embryos and subtracting this value from the total number of embryos collected. For larval cuticle preparation, terminally differentiated embryos were collected and dechorionated in bleach. After extensive washes in PBS-0.1% Triton X-100, embryos were mounted in Hoyer’s mountant/lactic acid (1:1). Samples were incubated overnight at 65°C. Imaging was performed on an Olympus BX-61, TRF microscope system.

### Culture and Live imaging of larval neuroblasts

Live imaging of isolated neuroblasts was performed using published methods (Pampalona et al., 2015). Brains were dissected from third instar larvae and then incubated in collagenase for 20 minutes. Brains were then manually dissociated with needles in fibrinogen (Sigma F-3879) dissolved in Schneider’s medium (SLS-04-351Q) on a 25mm glass bottom dish (WPI). Fibrinogen was then clotted by the addition of thrombin (Sigma T7513). Schneider’s medium was then added and a 5 µM slice at the centre of the cell was imaged every 2 minutes. Imaging was performed using a 100x objective (Oil, NA1.45) on a spinning disk confocal microscope. Data was processed using FIJI (Schindelin et al., 2012).

### Site-directed mutagenesis

An RNAi-resistant GFP-tagged Spindly transgene was generated by mutagenesis PCR targeting the shRNA-binding site (1611-caggacgcggttgatatcaaa-1632) at every third position of each codon, leaving unaltered the polypeptide sequence codified. The shRNA-resistant nucleotide sequence originated was 5’-caagatgccgtggacattaag-3’. A GFP-tagged variant of the full-length *spindly* cDNA cloned in a Gateway vector was kindly provided by Dr E. Griffis (School of Life Sciences, University of Dundee). This construct was used as a template in a site-directed mutagenesis PCR using the following mutagenic primers: Forward 5’-caagatgccgtggacattaagacggagttggaagctccagaattaattcc-3’, Reverse 5’-cttaatgtccacggcatcttgctgttccaagtttaagtcgatttctcg-3’ The DpnI_digested PCR products were used to transform E. coli and positive clones were sequenced to verify that only the appropriate mutation was present. The cDNA codifying for the RNAi-resistant GFP-Spindly was then subcloned in a pUASp K10 vector using KpnI-BamHI sites and used to generate transgenic animals. The same plasmid vector was used to generate Spindly mutant variants. Primers are available upon request.

### Protein extraction and Western blotting

0-3 hours-old embryos were collected on apple juice plates at different temperatures as indicated in the text. After dechorionation, embryos were lysed in RIPA Buffer (50 mM Tris-HCL pH 8.0, 150 mM NaCl, 1 % NP-40, 0.5 % Sodiumdeoxycholate, 0.1 % SDS) containing a mixture of protease inhibitors (Mini-complete EDTA-free-Roche, 1 tablet/10 ml). The homogenised tissue was incubated on ice for 30 minutes. Subsequently the samples were centrifuged twice at 13000 rpm for 30 minutes at 4°C to remove insoluble debris. Protein concentration was estimated by reading the absorbance at 600 nm using a photometer (Biophotometer, Eppendorf_Hamburg). For Western Blotting, 10 µm of proteins for each sample were heat-denatured, resolved on SDS-PAGE and transferred to PVDF membrane (Whatman, GE Healthcare) for immunoblotting. The membrane was immunoblotted with the following primary antibodies: rabbit-anti-Spindly 1:2000 (Griffis et al., 2007), rabbit-anti-actin 1:3000 (Sigma A2066), mouse-anti-tubulin 1:2000 (DSHB, 12G10). After extensive washes, membrane was incubated with HRP-conjugated antibodies: anti-rabbit-HRP 1:5000 (Thermo Scientific, #31460) and anti-mouse-HRP 1:2000 (Roche). Signals were detected using chemiluminescence on X-Ray film.

### Embryo immunostaining

Embryos for immunolabelling were collected every three hours at different temperatures as indicated in the text and dechorionated. Embryos were transferred in a 1:1 solution of Heptane and 4% formaldehyde in PBS for 20 minutes. Alternatively, embryos were incubated in a 1:1 solution of PEM Buffer containing 1 µM Taxol (Tocris) and Heptane to which 1 ml of 20% formaldehyde was added for 10 minutes. Taxol was added to stabilise microtubules and improve the staining of this cytoskeletal component (Karr and Alberts, 1986). After fixation, formaldehyde was replaced with methanol and vigorously shaken. After two washes in methanol, embryos were washed in PBS-0.1% Tween-20 (3 times, 15 minutes each) and incubated in blocking solution for 1 hour at room temperature. Primary antibodies were incubated overnight at 4°C, rotating. Primary antibodies were used at the following dilutions: rabbit anti-centrosomine 1:250 (kindly provided by Professor J. Raff), mouse anti-β-tubulin 1:50 (DSHB E7). Fluorophore-conjugated secondary antibodies (Invitrogen) were used at 1:250 and DAPI at 1 µg/ml. Specimens were mounted in MOWIOL/DABCO and imaged on Olympus BX-61, TRF fluorescence microscope. Images were processed using Volocity (Improvision, Perkin Elmer) and Adobe Photoshop.

### Ovary dissection and immunolabeling

1-3 old mated flies were placed on standard medium containing dry yeast for 24 hours at 25°C prior to dissection. Ovaries were dissected in phosphate buffer (PBS) and fixed in cold 4% formaldehyde in PBS for 15 minutes. Ovaries were washed repetitively in PBS- 0.1% Tween-20 and rotated for 2 hours in a solution of PBS- 0.1% Triton X-100 supplemented with 5% Foetal Calf Serum (FCS). F-actin staining was performed by adding fluorochrome-conjugated phalloidin (1:50) (Phalloidin-Alexa594, Invitrogen #A12381). Subsequently, samples were washed three times in PBS-0.1% Tween-20, 15 minutes each time on a rocking platform. Nuclei were stained with DAPI (1:1000) for 10 min. Ovaries were embedded in Mowiol/DABCO (Sigma Aldrich) and mounted on coverslips. Microscopy was performed using a confocal laser-scanning microscope (Leica SP2 and Leica TCS SP8). Images were processed using Volocity (Improvision, Perkin Elmer) and Adobe Photoshop. Image J (Abramoff et al., 2004) was used for image analysis.

### Egg chamber staging

Precise identification of egg-chamber stages was based on egg chamber morphology and stage-specific markers. At the transition between stage 8 and 9 of egg-chamber development, concomitantly to the initiation of border cell cluster migration, the anterior follicle cells start to migrate away from each other in an anterior to posterior direction to cover the oocyte. To unequivocally identify mid-stage 9 egg chambers we took advantage of the morphogenetic rearrangements of the follicular epithelium. Particularly a line was traced to connect the dorsal and ventral anterior-most cells in the follicular epithelium and the distance between the position of the follicle cells and the nurse cells (NC)-oocyte boundary was measured. For mid-stage 9 egg chambers an average distance value of 46 µm was calculated. Image analysis and distance measurements were performed on Image J.

### Quantification of the migration phenotypes

For scoring the migration of the border cell cluster, the egg chamber was divided in five regions: 0-25% f migration, 26-50% of migration, 51-75% of migration, 75-90% of migration and 100% complete migration. The distance travelled by the cluster was measured as distance between the posterior-most follicle cells and the leading border cell. This value was then normalised to the total distance, that is the distance between the posterior-most follicle cells and the NC-oocyte boundary and expressed as percentage. On the base of these measurements, the number of clusters falling in each region was counted and results represented in histograms. Image analysis and distance measurements were performed on Image J. Migration (MI) and Completion (CI) indices were calculated as previously described (Assaker at al. 2010). The MI is the measure of the mean distance travelled by the cluster. This index was calculated with the following formula:

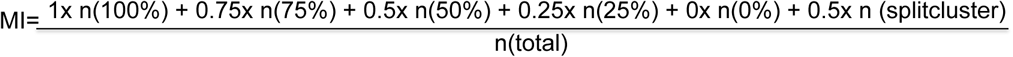

In the formula n (100%) represents the number of clusters that have completed migration, reaching the NC-oocyte border, n(75%) n50%) and n(25%) represent the number of clusters that have migrated 75, 50 and 25% of the total distance respectively. The formula includes also the number of dissociated clusters (n splitcluster). The CI index represents the number of clusters that have completed migration. CI is calculated as the ratio between the number of clusters that have migrated the total distance [n(100%)] and the total number of egg chambers examined [n(total)].

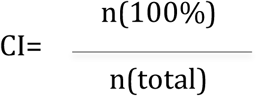

The indices were normalised to the wild-type control.

## Acknowledgements

We would like to thank Ryan Webster for his expert technical assistance and advice. We thank Jordan Raff (Oxford) for antibodies. We would like to thank the Developmental Studies Hybridoma Bank (DSHB, Iowa City, Iowa, USA) for monoclonal antibodies and the Bloomington (Indiana, USA) Drosophila Stock Centre for fly stocks.

## Author contributions

GDC, ERG, JJ, MRH and HAJM planned and designed the experiments. GDC and MRH performed all experiments. GDC, JJ, MRH and HAJM evaluated and interpreted the data. GDC, MRH and HAJM prepared the figures. GDC and HAJM wrote the manuscript and JJ edited the manuscript.

## Supplemental Material

### Movie1

Isolated neuroblast overexpressing UAS≫GFP::Spindly (red) and UAS≫mCherry::tubulin (cyan) under the control of *worniu*>Gal4. Corresponds to figure 1B. GFP::Spindly is recruited to the kinetochore shortly after NEB. At anaphase GFP::Spindly moves away from the metaphase plate towards the spindle pole. GFP::spindly remains at the spindle pole until mitosis is complete whereupon it delocalizes into the cytoplasm. Time stamp: hh:mm. Scale bar: 10µm.

### Movie2

Isolated neuroblast overexpressing UAS≫GFP::Spindly^ΔCt^ (red) and UAS≫mCherry::tubulin (cyan) under the control of *worniu*>Gal4. GFP::Spindly^ΔCt^ is never recruited to the kinetochore throughout mitosis and remains in the cytoplasm. Corresponds to Figure 1C. Time stamp: hh:mm. Scale bar: 10µm.

### Movie3

Isolated neuroblast overexpressing UAS≫GFP::Spindly^ΔSB^ (red) and UAS≫mCherry::tubulin (cyan) under the control of *worniu*>Gal4.GFP::Spindly^ΔSB^ localises similarly to the wild type, recruited to the kinetochore following NEB before stripping away from the metaphase plate towards the spindle pole where it remains until after mitosis. Corresponds to Figure 1D. Time stamp: hh:mm. Scale bar: 10µm.

### Movie4

Isolated neuroblast overexpressing UAS≫GFP::Spindly^ΔNt^ (red) and UAS≫mCherry::tubulin (cyan) under the control of *worniu*>Gal4. GFP::Spindly^ΔNt^ forms cytoplasmic puncta throughout interphase. In mitosis it is recruited to the kinetochore exhibiting dynamics similar to wild type, moving towards the spindle pole at anaphase. GFP::Spindly^ΔNt^ returns to cytoplasmic puncta following mitosis. Corresponds to Figure 1E. Time stamp: hh:mm. Scale bar: 10µm.

### Movie5

Isolated neuroblast overexpressing UAS≫GFP::Spindly^S34A^ (red) and UAS≫mCherry::tubulin (cyan) under the control of *worniu*>Gal4. GFP::Spindly^ΔS234A^ localises similarly to the wild type GFP::Spindly as well as GFP::Spindly^ΔSB^ localising to the kinetochore after NEB and moving towards the spindle pole upon anaphase onset where it remains until mitosis is complete. Corresponds to Figure 1F. Time stamp: hh:mm. Scale bar: 10µm.

